# Spontaneous recurrent seizures in an intra-amygdala kainate microinjection model of temporal lobe epilepsy are differentially sensitive to antiseizure drugs

**DOI:** 10.1101/2020.12.03.410266

**Authors:** Peter J. West, Kyle Thomson, Peggy Billingsley, Timothy Pruess, Carlos Rueda, Gerald W. Saunders, Misty D. Smith, Cameron S. Metcalf, Karen S. Wilcox

## Abstract

The discovery and development of novel antiseizure drugs (ASDs) that are effective in controlling pharmacoresistant spontaneous recurrent seizures (SRSs) continues to represent a significant unmet clinical need. The Epilepsy Therapy Screening Program (ETSP) has undertaken efforts to address this need by adopting animal models that represent the salient features of human pharmacoresistant epilepsy and employing these models for preclinical testing of investigational ASDs. One such model that has garnered increased interest in recent years is the mouse variant of the Intra-Amygdala Kainate (IAK) microinjection model of mesial temporal lobe epilepsy (MTLE). In establishing a version of this model, several methodological variables were evaluated for their effect(s) on pertinent quantitative endpoints. Although administration of a benzodiazepine 40 minutes after kainate (KA) induced status epilepticus (SE) is commonly used to improve survival, data presented here demonstrates similar outcomes (mortality, hippocampal damage, latency periods, and 90-day SRS natural history) between mice given midazolam and those that were not. Using a version of this model that did not interrupt SE with a benzodiazepine, a 90-day natural history study was performed and survival, latency periods, SRS frequencies and durations, and SRS clustering data were quantified. Finally, an important step towards model adoption is to assess the sensitivities or resistances of SRSs to a panel of approved and clinically used ASDs. Accordingly, the following ASDs were evaluated for their effects on SRSs in these mice: phenytoin (20 mg/kg, b.i.d.), carbamazepine (30 mg/kg, t.i.d.), valproate (240 mg/kg, t.i.d.), diazepam (4 mg/kg, b.i.d.), and phenobarbital (25 and 50 mg/kg, b.i.d.). Valproate, diazepam, and phenobarbital significantly attenuated SRS frequency relative to vehicle controls at doses devoid of observable adverse behavioral effects. Only diazepam significantly increased seizure freedom. Neither phenytoin nor carbamazepine significantly altered SRS frequency or freedom under these experimental conditions. These data demonstrate that SRSs in this IAK model of MTLE are pharmacoresistant to two representative sodium channel-inhibiting ASDs (phenytoin and carbamazepine) but not to GABA receptor modulating ASDs (diazepam and phenobarbital) or a mixed-mechanism ASD (valproate). Accordingly, this model is being incorporated into the NINDS-funded ETSP testing platform for treatment resistant epilepsy.

**Highlights:**

- An intra-amygdala kainate model of TLE was evaluated for pharmacoresistant seizures
- Administration of midazolam during status epilepticus did not affect mortality
- Model characteristics were evaluated over a 90-day natural history study
- Spontaneous seizures were resistant to phenytoin and carbamazepine
- Spontaneous seizures were sensitive to valproic acid, diazepam, and phenobarbital

## 1. Introduction

Pharmacoresistance has been defined by the International League Against Epilepsy (ILEA) as “failure of adequate trials of two tolerated, appropriately chosen and used antiepileptic drug schedules (whether as monotherapies or in combination) to achieve sustained seizure freedom” (Kwan et al., 2010). In accordance with this definition, it is widely recognized that nearly a third of all epilepsy patients experience spontaneous recurrent seizures (SRSs) that are refractory, or pharmacoresistant, to the present armamentarium of antiseizure drugs (ASDs). Therefore, the discovery and development of novel ASDs that are effective in controlling pharmacoresistant SRSs continues to represent a significant unmet clinical need. The epilepsy research community has acknowledged this need and codified its dedication to addressing this challenge within the NINDS epilepsy benchmarks. Specifically, benchmark area III highlights the need to develop new, or improve existing, ASDs for patients with refractory epilepsy (Long et al., 2016; Traynelis et al., 2020). As part of the greater epilepsy research community, and as recommended by three National Institute of Neurological Disorders and Stroke (NINDS) council working groups, the Epilepsy Therapy Screening Program (ETSP) has undertaken efforts to address this benchmark by adopting animal models that better represent the salient features of human pharmacoresistant epilepsy (Wilcox et al., 2019). One such model that has garnered increased interest and attention in recent years by a number of laboratories is the mouse variant of the Intra-Amygdala Kainate (IAK) microinjection model of mesial temporal lobe epilepsy (MTLE) (Iori et al., 2016; Jimenez-Pacheco et al., 2013; Kondratiuk et al., 2015; Li et al., 2008; Liu et al., 2013; Welzel et al., 2020)

Microinjection of the chemoconvulsant kainic acid (KA) into the rat amygdala to induce status epilepticus (SE) and the subsequent development of MTLE was first demonstrated in the late 1970s (Ben-Ari et al., 1980, 1979). These methods were successfully adapted for use in mice in the early 2000s (Araki et al., 2002; Shinoda et al., 2004). Unlike models that employ systemic injections of KA and lead to widespread damage, IAK microinjection in mice has been reported to cause limited and primarily unilateral hippocampal damage similar to human MTLE (Mouri et al., 2008). Furthermore, IAK mouse models have been reported to offer several advantages that make them particularly well suited for assessing the efficacy of antiseizure therapies (e.g. SRSs are reported to manifest after a short latent period and occur with a relatively stable frequency) (Jimenez-Pacheco et al., 2016). However, with the exception of carbamazepine (Iori et al., 2016; Welzel et al., 2020), the sensitivities or resistances of these SRSs to approved and commonly used ASDs is presently unknown.

The present study endeavored to establish and pharmacologically characterize an IAK microinjection model of MTLE under the auspices of the ETSP contract and as part of its ongoing efforts to internally validate and adopt improved animal models of human epilepsy with pharmacoresistant SRSs. Although most laboratories using this model have largely employed the methods established by Dr. Henshall and colleagues (Araki et al., 2002; Shinoda et al., 2004), several experimental variants with sometimes subtle methodological differences have been described (e.g. see (Welzel et al., 2020)). Furthermore, systematic testing of how altering several of these methodological variables affects outcomes (e.g., SE, neuronal damage, and SRS outcomes) has also been reported (Araki et al., 2002; Diviney et al., 2015; Silva et al., 2016). The study presented here, similarly, tested methodological variables, such as the effects of interrupting SE with midazolam, on neuronal damage and SRS outcomes. Subsequently, our version of the IAK mouse model of MTLE was evaluated by performing a 90-day natural history; outcomes including survival, latent period distribution, SRS frequencies and durations over 90 days, and SRS cluster analysis are described herein. Finally, SRSs in this model were evaluated in an initial study for their sensitivity to ASDs from several mechanistic classes. Drugs/doses tested were phenytoin (20 mg/kg, b.i.d.), carbamazepine (30 mg/kg, t.i.d.), valproate (240 mg/kg, t.i.d.), diazepam (4 mg/kg, b.i.d.), and phenobarbital (25 and 50 mg/kg b.i.d.). These data strongly suggest that SRSs resulting from IAK microinjection induced SE may be useful for identifying and differentiating novel therapies for pharmacoresistant epilepsy.

## 2. Materials and Methods

### 2.1. Animals

A total of 292 male C57BL/6J mice (4-5 weeks) from Jackson Laboratories (Bar Harbor, ME, U.S.A.) were used in these studies. All mice were group housed in a light- and temperature-controlled (12 h on/12 h off) environment and permitted access to food and water *ad libitum* before use. After surgery and for the remainder of the experiments, mice were single housed. All efforts were made to minimize the number and suffering of animals used. All experiments were performed in accordance with the NIH Guide for the Care and Use of Laboratory Animals and were approved by the University of Utah’s Institutional Animal Care and Use Committee. Principles outlined in the ARRIVE guidelines have been considered when planning experiments.

### 2.2. IAK mouse model of SE-induced epilepsy

Typically, 60 mice were surgically prepared per cohort. Buprenorphine was administered at 0.01-0.02 mg/kg, s.c., for the treatment of pain 1 hour before surgery. All surgical procedures were performed under general gas anesthesia (2–5% isoflurane in O2) and stereotaxic guidance. A Dremel drill was used to create three shallow screw holes where stainless steel anchor screws were affixed above the ipsilateral and contralateral frontal cortex and the contralateral somatosensory cortex. An additional hole was drilled over the right hemisphere, and a 22-gauge guide cannula was positioned on top of the dura mater (Figure 1A: from bregma: AP = -1.2 mm; ML = +3.3 mm). For the EEG depth electrode, holes were drilled over the right hemisphere dorsal hippocampus (Figure 1A: from Bregma: AP = -2.3 mm; ML = 2.5 mm) and cerebellum; a Teflon insulated platinum iridium depth electrode (MS333/6-B/SPC, Plastics One Inc. Roanoke, VA, U.S.A.) was then implanted into area CA1 (DV = -1.8 mm) and reference electrode over the cerebellum. Guide cannula, EEG electrodes, and a multipin socket were glued in place and a headcap was created with dental acrylic cement. Antibiotic ointment was applied around the incision site, a single dose of penicillin (60,000 units s.c.) was administered, and mice were allowed to recover in their individual home cages for ≥ 3 days.

**Figure 1:**
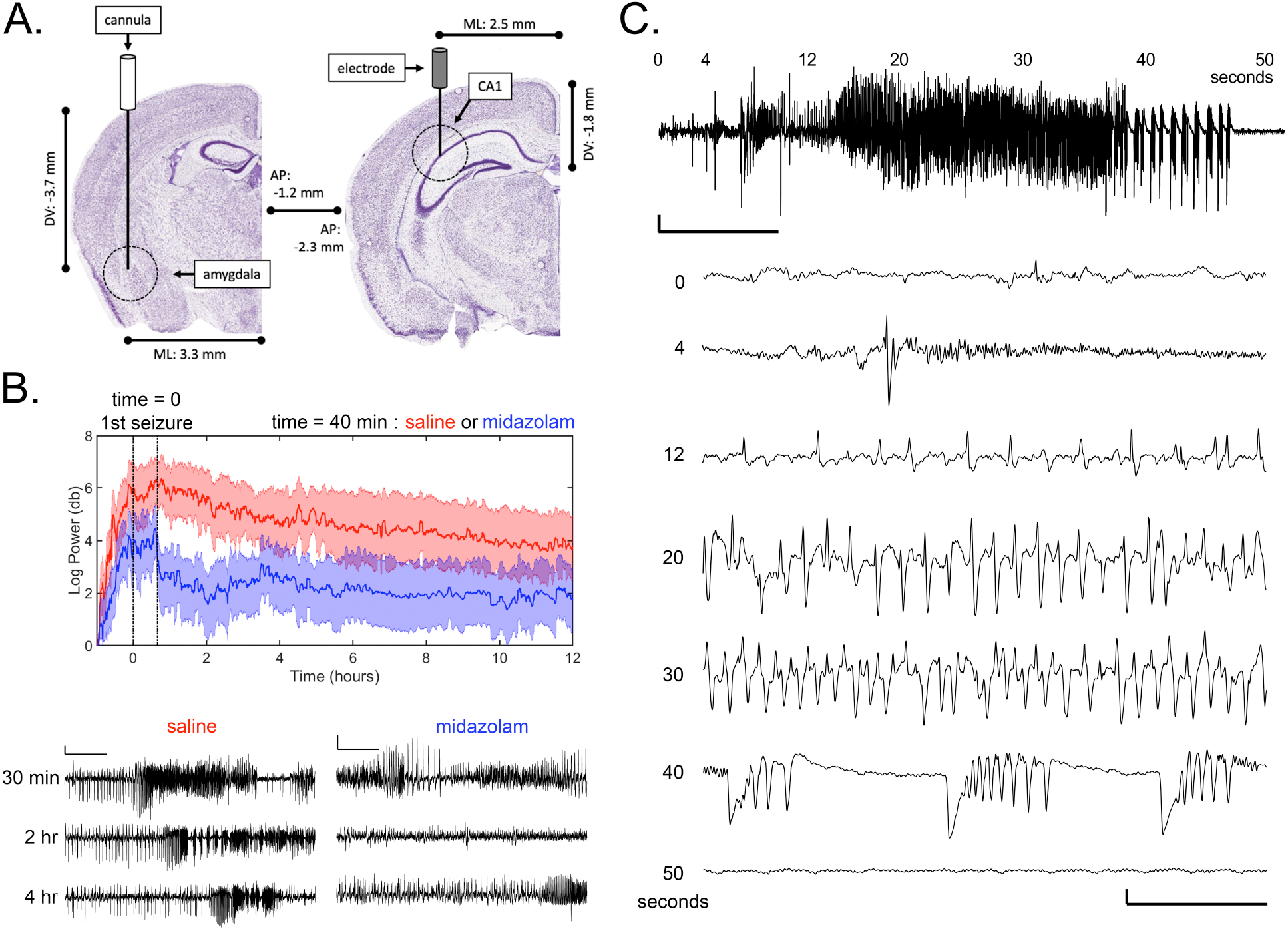
Microinjection of kainic acid (KA) into the mouse amygdala induces status epilepticus (SE) and the subsequent development of spontaneous recurrent seizures (SRS). A) Schematic showing coordinates for injection cannula and CA1 electrode implantation sites. Anatomic reference images courtesy of the Allen Mouse Brain Atlas (Lein et al., 2007). B) Gamma-band power resulting from microinjection of KA for mice given midazolam (N=9, Blue) or saline (N=9, Red) 40 minutes after detection of their first seizure. All mice EEG recordings were aligned to time = 0 when their first seizures were detected and time = 40 minutes when either 8 mg/kg midazolam or saline was administered. Data are represented by the mean (solid lines) and SEM (shaded area) for the Log10 of gamma frequency power (db). One-minute-long representative EEG traces for a saline-treated (left) and midazolam-treated (right) IAK mouse are shown below. Relative to time zero, traces were captured at 30 minutes (before saline or midazolam treatment), 2 hours (approximate time of largest midazolam effect), and 4 hours (after recovery from midazolam). Note the attenuation (at 2 hr) and recovery (at 4 hr) of large amplitude EEG spikes in the midazolam group. Scale bar represents 2 mV and 10 seconds. C) Representative EEG seizure recorded from an IAK mouse with an ipsilateral CA1 hippocampal depth electrode during a Racine stage 4-5 behavioral seizure. Scale bar represents 2 mV and 10 seconds. Time points (seconds) above the trace correspond to the expanded traces below (total duration = 2 seconds; bottom scale bar represents 2 mV and 0.5 seconds). Similar to EEG waveforms documented in other seizure models (e.g. see (Williams et al., 2009)), seizure initiation begins with a large EEG spike (4 sec), followed by progressive high amplitude high frequency EEG spikes (12, 20, and 30 sec), large amplitude waves containing multiple EEG spikes (40 sec), and finally post-ictal depression (50 sec).

While the majority of published experiments using variants of this model report injecting 0.3 µg of KA in a volume of 0.2 µL directly into the basolateral amygdala (BLA), some experimentation with alternate KA doses has been noted (Welzel et al., 2020). Also, KA storage conditions and lot-to-lot differences were considered for their contributions to inter-experiment variability. Accordingly, our early pilot experiments assessed the impact of varying doses of Kainic Acid Monohydrate (KA, 0222, Tocris, Minneapolis, MN) on outcomes. These pilot studies were conducted after combining several lots of KA, creating a single stock solution (1.5 mg/mL in 0.9% saline), and freezing aliquots at -20°C. Dose was adjusted by increasing or decreasing the volume of the stock solution injected (0.2 – 0.3 µL). For this particular stock solution of KA, these pilot experiments (data not shown) demonstrated that favorable outcomes (low mortality and high percentages of mice developing SRSs) were achieved after injection of 0.39 µg in 0.26 µL saline. Therefore, this dose was chosen for all of the experiments described.

On the day of KA injection, a single 1.5 mg/mL KA aliquot was thawed and placed in a glass syringe pump (Pump 11 Elite, Harvard Apparatus, Holliston, MA, U.S.A.). In conscious mice under mild restraint, the attached injection cannula (c313I/SPC, Plastics One Inc. Roanoke, VA, U.S.A.) was inserted into the guide cannula and slowly lowered into the BLA at a depth of 3.7 mm below the dura (Figure 1A). KA (0.39 µg in 0.26 µL) was then injected at a rate of 50 nL/second. The injection cannula was left in place for an additional 2 min after KA microinjection to allow diffusion into the brain tissue and minimize movement of KA up the cannula track. Status Epilepticus (SE), as defined by continuous EEG spikes >1 Hz and concomitant stage 3-5 seizures, typically began approximately 30 minutes after KA microinjection (Figure 1B). In experiments where SE was interrupted with a benzodiazepine, midazolam (8 mg/kg, i.p.) was administered 40 min after initiation of SE (Figure 1B). For the 18 mice used to study SE (Figure 1B), continuous EEG recordings were made for an additional 12 hours. For all other experiments, mice were detached from their tethered EEG cables 40 minutes after confirmation of SE, given lactated ringers (0.25 mL, i.p.), and allowed to recover for 3 days in their individual home cages where behavioral health (e.g., regular eating, drinking, and grooming) were monitored. This approach was adopted because, in pilot experiments, it was noted that recovery from SE was improved when mice were allowed to freely move about their cages and care for themselves without the hindrances caused by an EEG tether.

### 2.3. Tissue processing, FluoroJade-B staining, and Imaging

Whole brains were removed and allowed to post-fix in 4% PFA overnight at 4 °C before being rinsed with PBS and transferred to a 30% sucrose solution for overnight cryoprotection. Thirty micrometer thick coronal sections were sliced on a freezing stage microtome and stored at 4 °C in PBS for staining. FluoroJade-B (FJB) staining was performed using 3 hippocampal slices per animal and batch processed. Briefly, sections were mounted on glass slides, dried, and immersed in a descending ethanol series (100% for 3 min, and 90%, 70%, and dIH_2_O for 1 min each). Slices were blocked in 0.06% KMnO_4_ for 30 min, followed by a 2 min rinse in dIH_2_O, and then incubated in 0.001% FJB solution for 15 min. Slides were rinsed 5 times with dIH_2_O for 1 min each, dehydrated in an ascending ethanol series (70%, 90%, and 100% for 3 min each), and coverslipped with DPX mounting media. Digital images were obtained with a Zeiss Axioskop microscope equipped with an AxioCam digital camera system and AxioVision 3.0 software (Zeiss, Oberkochen, Germany). Three hippocampal subregions (CA1, CA3, and Hilus, both ipsilateral and contralateral to the KA injection) were scored for positive or negative FJB staining by two independent reviewers who were blinded to treatment group.

### 2.4. Video-EEG monitoring and assessment of SRSs

Three days after KA microinjection induced SE, IAK mice were connected to their EEG electrode assembly via a tether to a rotating commutator (8BSL3CX, Plastics One, Inc, Roanoke, VA, U.S.A.), which in-turn was connected to an EEG100C amplifier and digitized by an MP160 Recording system (BioPac Systems Inc., Goleta, CA, U.S.A.). Simultaneously, video was captured by a DVP 7020BE Capture card (Advantech, Milpitas, CA, U.S.A.). Video and EEG data were synchronized and written to disk using our custom software (Thomson and White, 2014). EEG and video data were continuously recorded from mice in this fashion, 24 hours per day and 7 days per week, for up to 90 days. EEG and video data were manually analyzed in a blinded and randomized fashion on a daily basis. Seizures were defined as high frequency (>5 Hz) and high amplitude (>2x baseline) poly-spike discharges of >5 seconds duration (Mouri et al., 2008; Pitkänen et al., 2005), and seizure termination was defined as a return of amplitude and frequency to baseline values with or without postictal depression. Behavioral seizures were scored as non-convulsive, focal Racine stage 1-3 (facial automatisms or unilateral forelimb clonus) or secondarily generalized Racine stage 4-5 (bilateral forelimb clonus, rearing, and/or falling).

### 2.5. Treatment with ASDs

All ASDs tested in these studies dissolved in saline (0.9% NaCl) with the exception of CBZ, which was dissolved in 0.5% methylcellulose (Sigma). Doses were chosen based on efficacy and toxicity results in pilot studies or from previously reported data in rats from established models including the LTG-kindled rat (LTG-R) model and the i.p. KA spontaneously seizing rat model of TLE (Metcalf et al., 2019; Thomson et al., 2020). In all six experiments, separate cohorts of IAK mice were generated as described. After 3-4 weeks, only mice that had experienced at least one SRS were enrolled in these studies. The five ASDs examined in these experiments were phenytoin (5,5-Diphenylhydantoin sodium salt, Sigma D4505, PHT, 20 mg/kg b.i.d.), carbamazepine (Sigma C8981, CBZ, 30 mg/kg t.i.d.), valproic acid (Valproic acid sodium salt, Sigma P4543, VPA, 240 mg/kg t.i.d.), diazepam (Sigma D0899, DZP, 4 mg/kg b.i.d.), and phenobarbital (Sigma P1636, PB, 25 and 50 mg/kg b.i.d.). All ASDs were administered via intraperitoneal (IP) injection. A sub-chronic dosing “cross-over” experimental design was chosen for these experiments (Thomson et al., 2020). Briefly, after a 7-day baseline observation period, half of the mice received ASD and the other half received vehicle for 5 days. Following a 2-day washout period where no drug or vehicle was administered, each group was administered the opposite treatment for an additional 5 days. Finally, all mice were observed for SRSs without treatment for an additional 7 days in order to monitor any reversal of ASD-induced effects.

### 2.6. Data Analysis and Statistics

Post-KA injection SE was analyzed by first applying a 60 Hz notch filter and then quantifying gamma frequency (20-70 Hz) power of the EEG recording as previously described (Lehmkuhle et al., 2009). This filtering approach was used to reduce motion artifact and other sources of noise in the electrophysiological recording. For all other experiments, the total number of SRSs were quantified per IAK mouse and subsequently averaged for all mice in each study on a daily or weekly basis. A representative SRS EEG recording can be seen in Figure 1C, and an additional SRS with an associated video recording of the mouse behavioral seizure can be seen in Video 1. Seizure freedom was defined as zero seizures occurring between the first and last dose of drug. SRSs clusters were defined, according to Lim et al., as the occurrence of one or more seizures per day for at least three consecutive days and at least five seizures within the cluster period (Lim et al., 2018). All numerical values were expressed as the mean ± SEM, and all error bars on graphs represent SEM. Statistical differences between categorical data (general outcomes, FJB + and –, and seizure freedom percentages) were compared using a Fisher’s Exact Test. Differences in latent periods between Midazolam and No Midazolam groups were compared using the Mann-Whitney Test. Differences in average weekly SRS frequencies over 90 days between Midazolam and No Midazolam groups were compared with a two-way repeated-measures ANOVA. Differences in average weekly SRS frequencies between ASD and vehicle groups were compared with the Wilcoxon Matched-Pairs Signed Rank Test. All statistical analyses were performed using GraphPad Prism V8.4.3 (GraphPad Software, San Diego, CA); *, **, ***, or **** indicate a p-value of <0.05, <0.01, <0.001, and <0.0001, respectively.

## 3. Results

### 3.1. Assessment of benzodiazepine treatment during SE on SRS outcomes and hippocampal damage

As originally described (Araki et al., 2002; Shinoda et al., 2004), the IAK mouse is a model of acquired TLE where the acquired insult, SE, is introduced by microinjection of the chemoconvulsant KA directly into the BLA. In subsequent years, subtle methodological variants of this model have been adopted by a number of laboratories (Iori et al., 2016; Jimenez-Pacheco et al., 2013; Kondratiuk et al., 2015; Li et al., 2008; Liu et al., 2013; Welzel et al., 2020). In order to establish a version of this model in our laboratory, we began by directly examining the effects of these methodological variables. Since many variants of this animal model administer a benzodiazepine 40 minutes after initiation of SE to “minimize morbidity and mortality” (Diviney et al., 2015), we examined the effects of administering the benzodiazepine midazolam (8 mg/kg, i.p., 40 minutes after initiation of SE) on a number of SRS outcomes and hippocampal damage. The immediate effects of midazolam (N=9) or saline (N=9) injections on SE are illustrated in Figure 1B. The largest observed midazolam-mediated attenuation of gamma band EEG power occurred 1-2 hours after initiation of SE; although there was a noticeable trend, this attenuation was not significant relative to saline injected controls (p = 0.0524, Mann-Whitney test).

Next, we directly tested the effects of administering midazolam (8 mg/kg, i.p., 40 minutes after initiation of SE) on mortality and other endpoints important to our planned pharmacology experiments. Two cohorts of IAK mice were generated: those receiving injections of midazolam during SE (N=27), and those receiving saline control injections (N=24). These mice were monitored for SRSs using 24/7 video EEG for 90 days. Figure 2A illustrates the outcomes of this experiment as percentages of mice in three categories: those that survive for 90 days and develop SRSs, those that did not develop SRSs, or mice that died at any timepoint over 90 days. No differences were observed in the percentages between the two cohorts (P > 0.9999, Fisher’s Exact Test) indicating that administering midazolam did not affect these outcomes.

**Figure 2:**
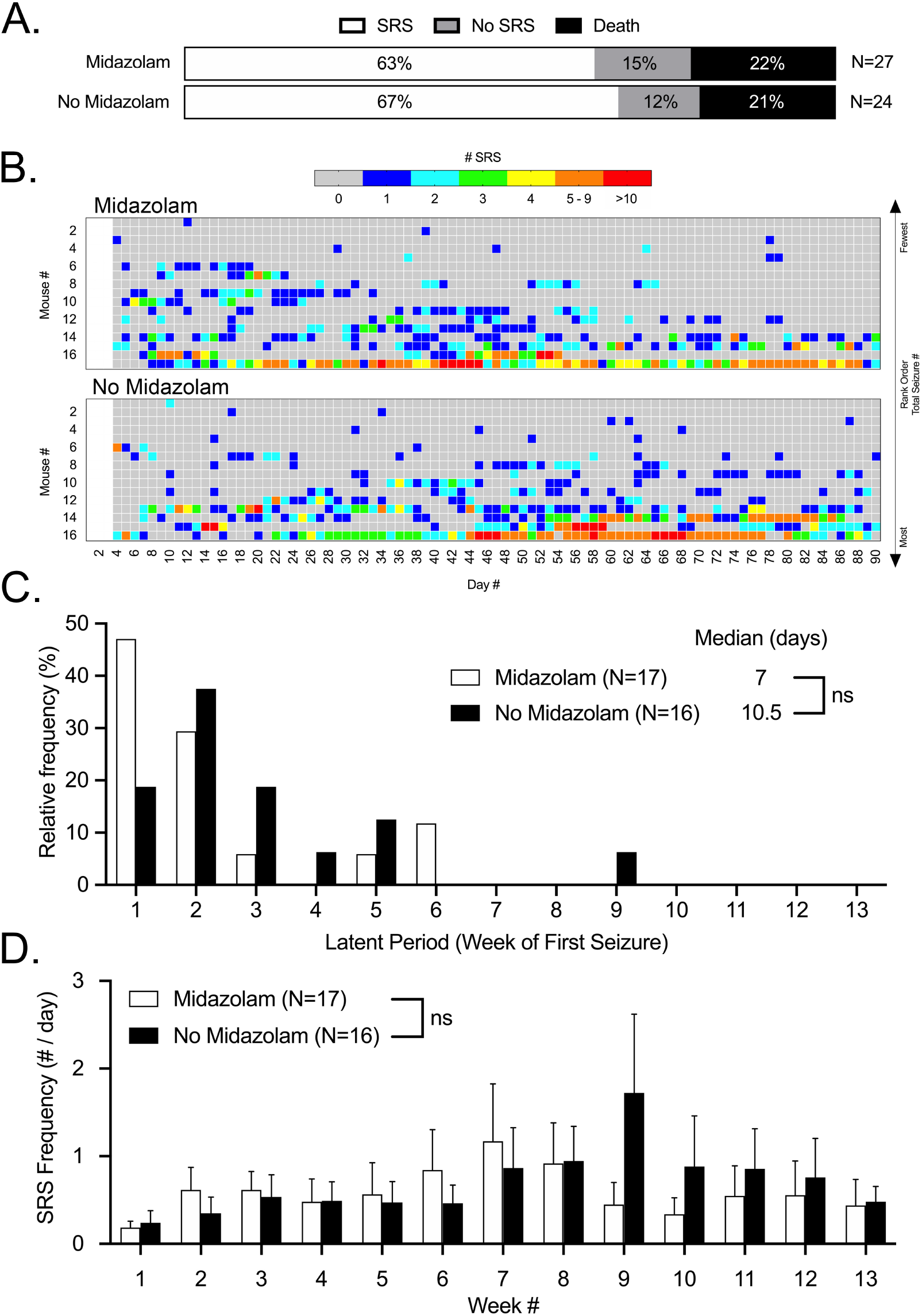
Midazolam treatment during SE does not significantly affect survival or the development of epilepsy in IAK mice. A) Percentages of those animals developing spontaneous recurrent seizures (SRS), not developing SRS, and died in a cohort of mice that received 8 mg/kg midazolam 40 minutes after initiation of SE and another cohort of mice that did not receive midazolam. B) Heatmaps depicting the daily number of SRSs experienced by each mouse in both cohorts over a 90-day video-EEG observation period. From top to bottom, mice were rank ordered by the total number of SRSs experienced over 90 days from lowest to highest respectively. Note: video EEG was not recorded for the first 3 days after SE. C) Relative frequency distributions for latent periods and group medians. Open bars represent midazolam treated mice and closed bars represent mice whose SE was uninterrupted. Median latent periods were not significantly different (ns, Mann-Whitney test). D) Weekly averages for SRS daily frequencies over the 90-day natural history observation period. Open bars represent midazolam treated mice and closed bars represent mice whose SE was uninterrupted. There were no significant differences in seizure frequency between these cohorts over the entire period (ns, two-way repeated-measures ANOVA).

In those mice that went on to develop SRSs and survived the entire 90 days of the video EEG observational period, we also compared latency to the first SRS (latent period) and average weekly SRS frequency between mice receiving midazolam (N=17) and those that did not (N=16). Heatmaps of the daily number of SRSs experienced by each mouse in this study are illustrated in Figure 2B. A frequency distribution histogram of latent periods, binned by week, is illustrated in Figure 2C. The median latent periods for mice receiving midazolam and those that did not, 7 and 10.5 days respectively, were not significantly different (P = 0.1584, Mann-Whitney test). Furthermore, as shown in Figure 2D, there were no significant differences in daily SRS frequencies, averaged weekly, between the midazolam-treated and untreated cohorts (F(1, 403) = 0.4967, P = 0.4814, two-way repeated-measures ANOVA).

Finally, in separate cohorts of midazolam-treated (N=10) and untreated (N=9) IAK mice, we examined hippocampal damage 48 hours after SE by qualitatively evaluating tissue sections of the dorsal hippocampus that were processed for FJB. Three hippocampal subregions (CA1, CA3, and Hilus, both ipsilateral and contralateral to the KA injection) were scored for positive or negative FJB staining by two independent reviewers who were blinded to treatment group. The results of this analysis are illustrated in Figure 3. Figure 3A shows representative images from two different mice for each experimental group; these representative images illustrate the range of results we observed, from those with limited ipsilateral CA3 damage to those that showed broad and clear FJB+ cells across CA3, CA1, and the hilus area of the dentate gyrus, both ipsilaterally and contralaterally. For both midazolam-treated and -untreated mice, 100% of their tissue sections had prominent FJB+ staining in the ipsilateral CA3 and 0% of these same sections were FJB+ in the contralateral hilus. Regarding the remainder of the evaluated subregions, although there was a trend towards a higher percentage of mice demonstrating FJB+ classifications in the no-midazolam group, these differences were not statistically significant (P values between 0.1409 – 0.5820, Fisher’s exact test).

**Figure 3:**
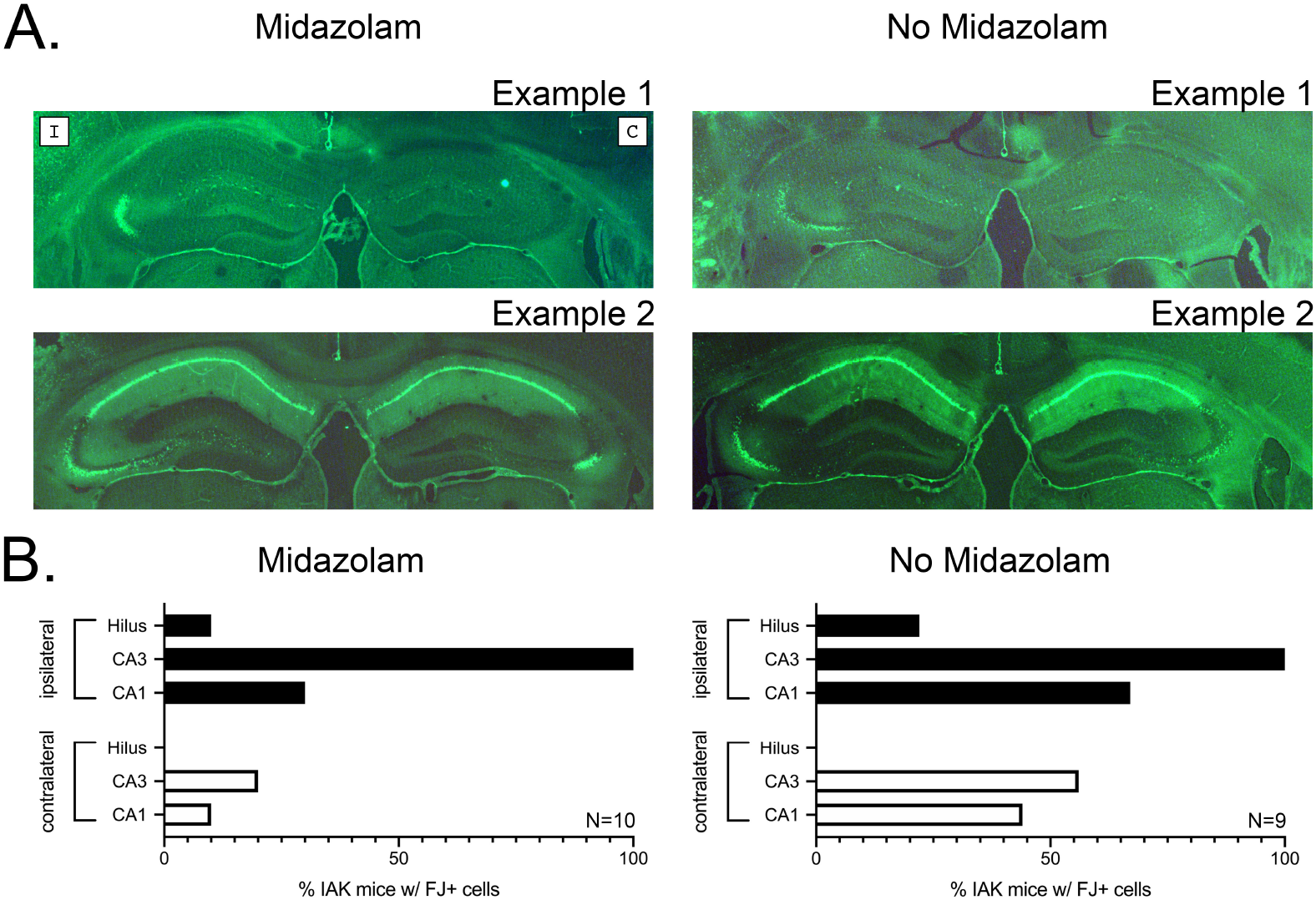
Midazolam treatment during SE does not significantly affect hippocampal damage 48 hours after SE. A) Representative photomicrographs of FluoroJade-B stained neurons in the dorsal hippocampi of midazolam-treated (Left) and non-treated (Right) IAK mice. As depicted in Example 1 of the Midazolam images, the left hemisphere is ipsilateral “I” to the site of KA injection, and the right hemisphere is contralateral “C”; this orientation applies to all images. In both midazolam and no midazolam images, Example 1 represents isolated ipsilateral CA3 damage and Example 2 represents extensive ipsilateral and contralateral damage. B) Quantified percentages of mice with FluoroJade-B positive cells in each subdivision of the ipsilateral and contralateral dorsal hippocampus reveals no significant differences between midazolam-treated and non-treated groups. In both graphs, solid bars represent ipsilateral percentages and open bars represent contralateral percentages.

Since injection of midazolam 40 minutes after the start of SE did not significantly affect hippocampal damage, improve mortality, significantly affect latent periods, or significantly affect SRS frequencies, we elected to omit benzodiazepine administration from all subsequent experiments.

### 3.2. Development of SRSs in this intra-amygdala kainate (IAK) microinjection model of temporal lobe epilepsy (TLE)

Before proceeding to an examination of the sensitivity of SRSs to antiseizure drugs (ASDs) in this variant of the IAK mouse model, we performed a comprehensive natural history experiment in order to characterize outcomes, survival timelines, latent periods, SRS frequencies, behavioral seizure scores, SRS durations, and SRS clustering under our experimental conditions. The results of this natural history study are illustrated in Figures 4 and 5. In addition to the 24 “no-midazolam” IAK mice that were described in Figure 2, an additional 77 mice were included in this study; thus, the total number of IAK mice examined under these experimental conditions and included in this study was 101. Figure 4A illustrates the general outcomes observed over this 90-day experiment: 54% of these mice developed SRSs and survived for the entire 90 days of the experiment, 10% did not develop SRSs, and 36% died at varying timepoints. Of the 36% that died, 50% of these mice developed SRSs before they died and were not included in the “SRS” group that developed seizures *and* survived the entire 90 days; had these mice been included, the total percentage of mice that developed SRSs in this study was 72%. The survival curve for these IAK mice is illustrated in Figure 4B. In total, 20% of IAK mice died within the first 21 days, and an additional 15% died over the remaining 69 days of the experiment. The cause of death was not determined.

**Figure 4:**
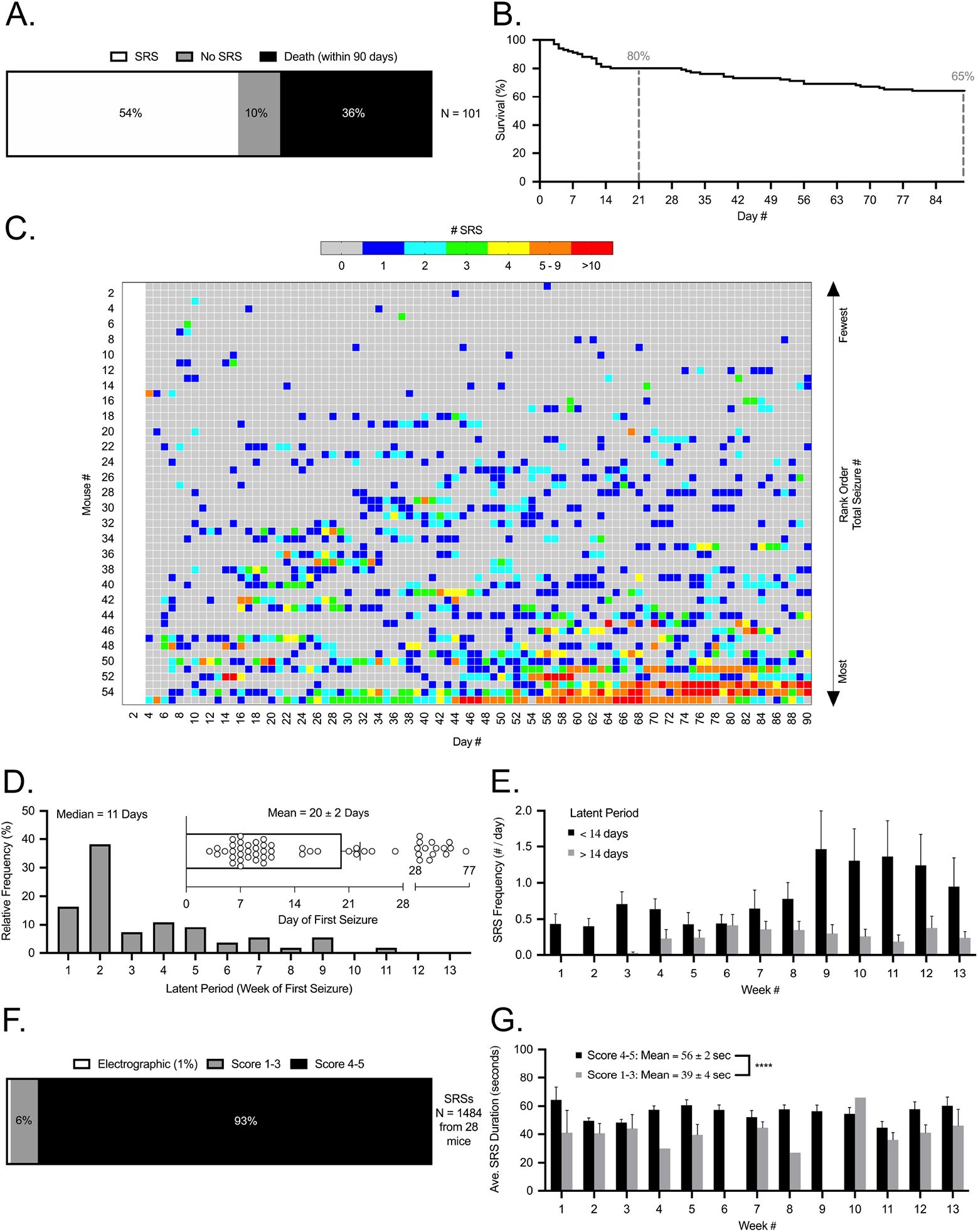
Natural history and characterization of SRSs in this IAK microinjection model of MTLE. A) Percentages of SRS, no SRS, and death for a cohort of 101 IAK mice. B) Survival curve for the 36 mice that died at varying times over the course of the 90-day observation period. Dashed lines highlight 80% survival at 21 days and 65% survival at 90 days. C) Heatmaps depicting the daily number of SRSs experience by each mouse over the 90-day video-EEG observation period. From top to bottom, mice were rank ordered by the total number of SRSs experienced over 90 days from lowest to highest respectively. Note: video EEG was not recorded for the first 3 days after SE. D) Relative frequency distribution of latent periods for all mice in this study. Median latent period was 11 days (Mean latent period was 20 ± 2 days). E) Weekly averages for SRS daily frequencies over the 90-day natural history observation period. Solid bars represent 30 mice whose latent period was <14 days, and gray bars represent 25 mice whose latent periods were >14 days. F) Percentages of SRSs that were accompanied by Racine score 4-5 behavioral seizures (93%), score 1-3 behavioral seizures (6%), or without obvious behaviors (1%). G) Weekly averages for SRS durations over the 90-day observation period. Solid bars represent seizure durations for score 4-5 behavioral seizures, and gray bars represent seizure durations for score 1-3 behavioral seizures. For panels F and G, a subset of 1484 SRSs were analyzed from a total of 28 mice.

**Figure 5:**
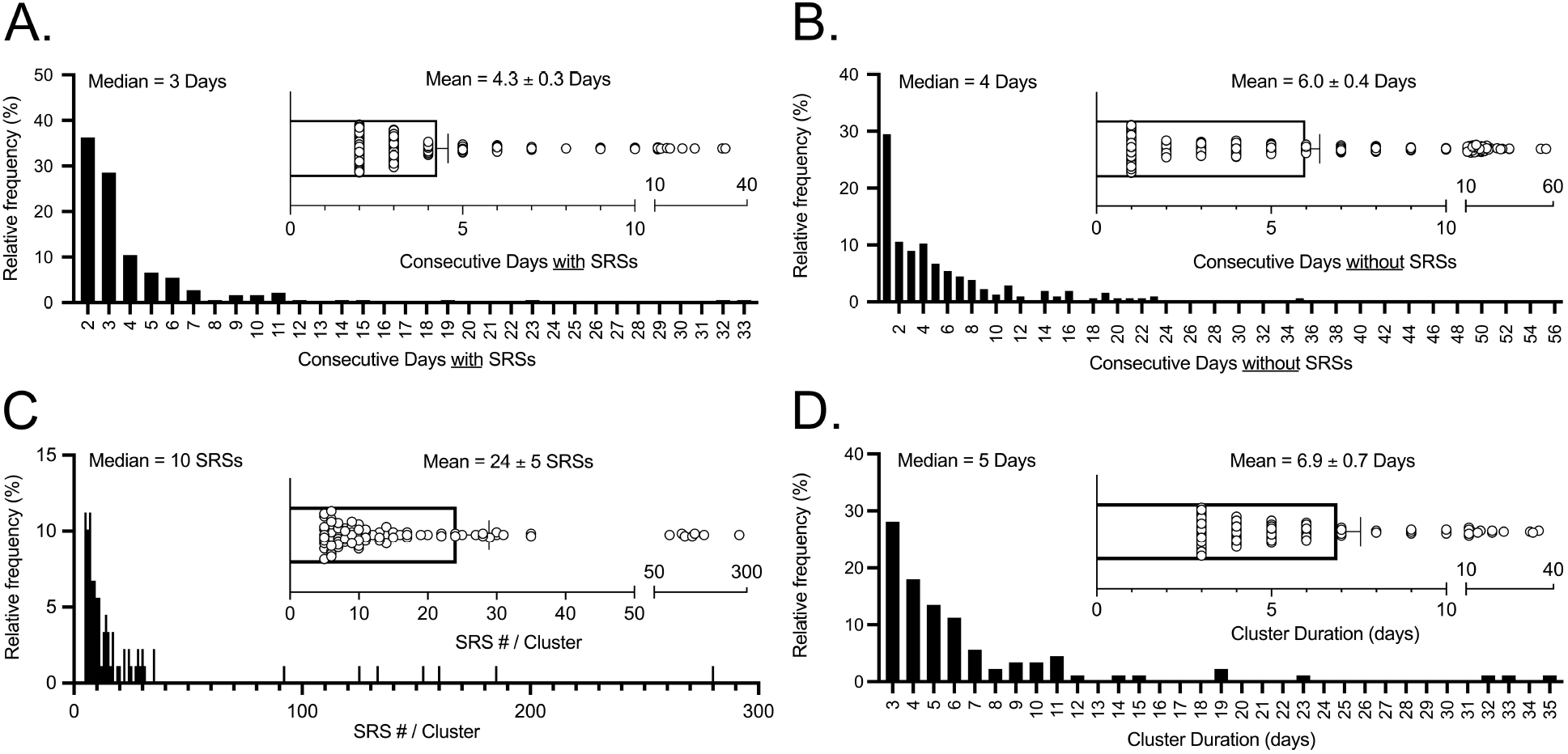
Natural history and characterization of SRS clusters in this IAK microinjection model of MTLE. A) Relative frequency distribution of the number of consecutive days IAK mice experienced SRSs. Median was 3 days. Inset bar graph represents mean (4.3 ± 0.3 days) of this distribution and individual consecutive day events. B) Relative frequency distribution of the number of consecutive days IAK mice did not experienced SRSs. Median was 4 days. Inset bar graph represents mean (6.0 ± 0.4 days) of this distribution and individual consecutive day zero-SRSs periods. C) Relative frequency distribution of the total number of SRSs per seizure cluster. Median was 10 SRSs. Inset bar graph represents mean (24 ± 5 SRSs) of this distribution and individual SRS number per cluster. D) Relative frequency distribution of seizure cluster durations (days). Median was 5 days. Inset bar graph represents mean (6.9 ± 0.7 days) of this distribution and individual seizure cluster durations.

Heatmaps of the daily number of SRSs experienced by the 55 mice that developed SRSs and survived 90 days in this study are illustrated in Figure 4C. The majority of these SRSs were secondarily generalized behavioral seizures as observed by others (Iori et al., 2016; Jimenez-Pacheco et al., 2013; Kondratiuk et al., 2015; Li et al., 2008; Liu et al., 2013; Welzel et al., 2020). A frequency distribution histogram of latent periods, binned by week, is illustrated in Figure 4D. The median latent period for these mice was 11 days. The total number of SRSs experienced by individual IAK mice each week were used to calculate average daily SRS frequencies (# / day); these average daily SRS frequencies, averaged across each IAK mouse for each week of this experiment, are plotted in Figure 4E. As illustrated, average daily SRS frequencies progressively increased from week 1 through 8 and achieved a maximum of 1.5 ± 0.5 SRSs / day on week 9 in the 30 mice that had latent periods less than 14 days. As illustrated in Figure 4F, 99% of EEG SRS events were accompanied by behaviors consistent with focal seizures (Racine score 1-3: 6%) or secondarily generalized tonic-clonic seizures (Racine score 4-5: 93%). The average durations of these seizures were 39 ± 4 seconds for score 1-3 seizures and 56 ± 2 seconds for score 4-5 seizures; this difference was significant (p < 0.0001, two-tailed student t test). These seizure durations did not appreciably change from week 1 through 13 of this natural history study.

Rather than having a regular occurrence, SRSs in these IAK mice often happened in closely grouped series separated by a number of seizure-free days. To better understand these SRS clusters, we quantified the relative frequencies of consecutive days with or without SRSs (Figures 5A and 5B respectively). For each IAK mouse, we defined “single days” as ≥1 SRS on a given day that was preceded and followed by an SRS-free day. Consecutive SRS days were defined as ≥1 SRS per day that were uninterrupted for ≥2 days. Finally, consecutive SRS-free days were defined as ≥1 day without SRSs. Using these definitions, we found that 177 of 359 SRS events (49%) were classified as “single day” events, and the remaining 182 events (51%) were strings of consecutive seizure days. The range of consecutive days with SRSs was 2 to 33 days with a median of 3 days and a mean of 4.3 ± 0.3 days (Figure 5A). Ninety percent (90%) of these consecutive day SRS events lasted 7.7 days or fewer. As for consecutive days without SRSs (Figure 5B), 90% of these occurrences lasted 15 days or fewer. The median number of consecutive days without SRSs was 4 days and the mean was 6.0 ± 0.4 days.

Defining seizure clusters by consecutive days with or without SRSs is one of many possible ways of categorizing and quantifying these events. In fact, numerous other definitions of seizure clusters have been proposed and used (Bortel et al., 2010; Goffin et al., 2007; Hosford et al., 2016; Lim et al., 2018; Williams et al., 2009). According to Lim et al. (2018), seizure clusters are defined as the occurrence of one or more seizures per day for at least three consecutive days and at least five seizures within the cluster period. Using this definition, we quantified the number of SRSs per cluster and cluster durations. The frequency distribution of SRS number per cluster is illustrated in Figure 5C; the median of this distribution was 10 SRSs and the mean was 24 ± 5 SRSs. Finally, the relative frequency distribution for cluster duration is illustrated in Figure 5D; the median of this distribution was 5 days and the mean was 6.9 ± 0.7 days. These SRS clustering characteristics were considered when interpreting ASD effects in the following experiments.

### 3.3. Pharmacological sensitivity of SRSs

With only a few exceptions (Iori et al., 2016; Welzel et al., 2020), the sensitivities or resistances of SRSs to existing ASDs in the IAK mouse model has not been tested. In order to assess if this model of TLE results in pharmacoresistant SRSs, we tested five ASDs with varying mechanisms of action for their abilities to affect SRS frequency or seizure freedom in our version of the IAK mouse.

#### 3.3.1. Phenytoin

Figure 6 (panels A-D) illustrates the effect of a sodium channel inhibiting ASD, PHT, on SRSs under our experimental conditions. Bar graphs in panels A-B represent the average daily number of SRSs over the course of the 26-day PHT experiment. Baseline and washout data (white bars) are the average for all mice in these experiments and, because of the cross-over experimental design employed, are the same for both the vehicle graph (Figure 6A) and the PHT graph (Figure 6B). Finally, all daily SRS data from each segment of the cross-over were averaged (gray or black bars, days 8-12 and 15-19). The average number of daily SRSs per mouse over the baseline period ranged from 0.7±0.4 to 2.2±0.8 SRSs/day (Figure 6, panels A-B); the weekly average was 1.6±0.4 SRSs/day (Figure 6C, baseline). There was no significant difference between the VEH (1.3±0.4 SRSs/day) and PHT (1.3±0.6 SRSs/day) 5-day treatment averages (P = 0.7251, Wilcoxon Test, Figure 6C). Finally, as illustrated in Figure 6D, statistical differences in the number of mice that were seizure-free during their treatment periods were computed using the Fisher’s Exact test. Five of 21 mice were seizure-free during VEH treatment and 7 of 21 during PHT treatment; this difference was not significant (P = 0.7337).

**Figure 6:**
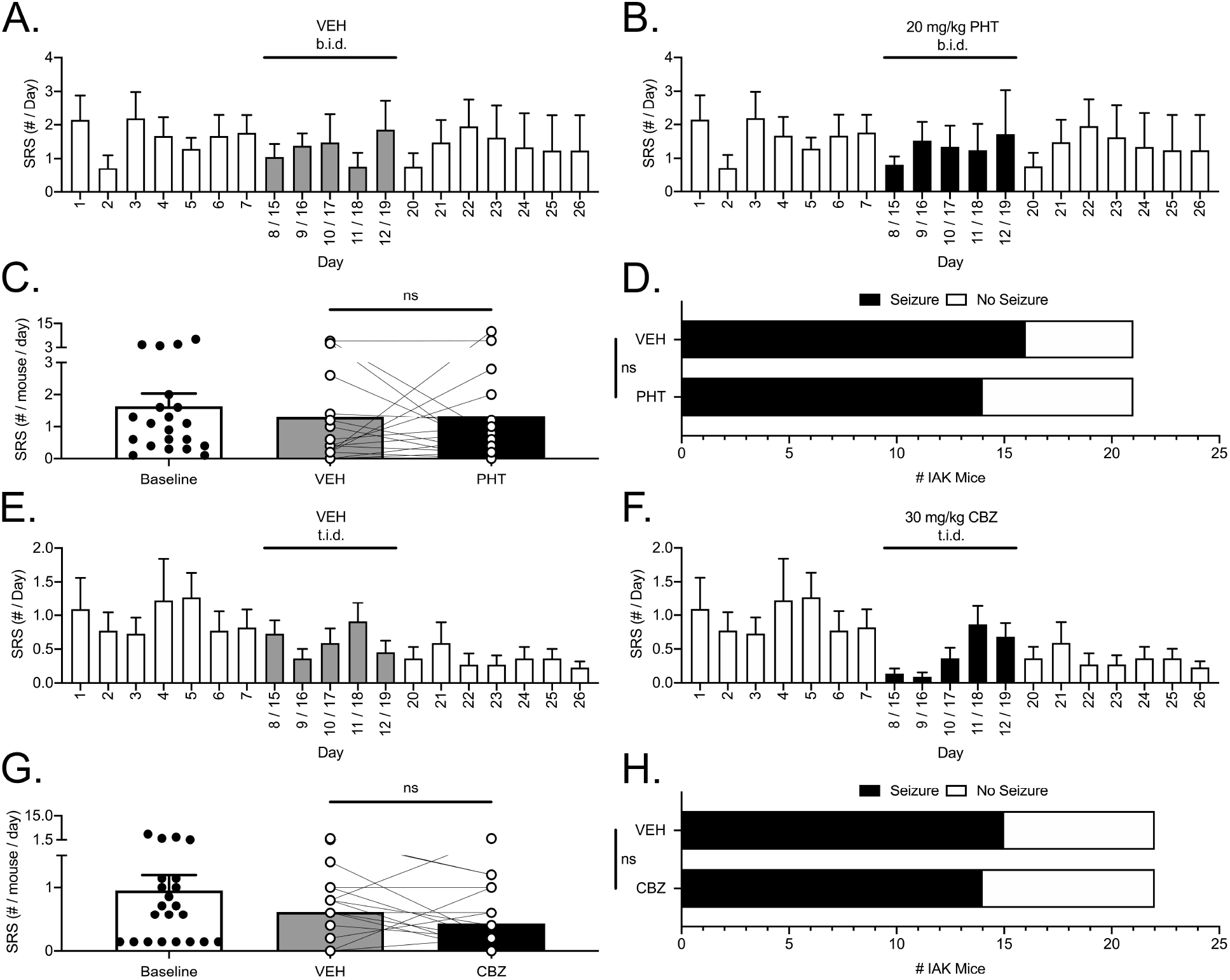
SRSs in IAK mice are resistant to phenytoin (40 mg/kg/day) and carbamazepine (90 mg/kg/day). A-B and E-F) Daily average SRS frequencies over 26-day evaluation periods for vehicle (A: saline or E: methylcellulose) and (B) PHT or (F) CBZ treatments. In all panels, open bars represent baseline and washout periods, while gray bars represent vehicle treatment averages (VEH) and solid black bars represent PHT or CBZ treatment averages for both halves of the crossover. C and G) SRS frequency daily averages, and individual mouse data points, for baseline (open bar), vehicle (gray bar), and (C) PHT or (G) CBZ (black bar) treatments. Vehicle and PHT data were not significantly different (ns, Wilcoxon Test). Likewise, Vehicle and CBZ data were not significantly different (ns, Wilcoxon Test). D and H) Comparisons of the percentages of SRS-free mice during vehicle and (D) PHT or (H) CBZ treatments. Solid black bars represent percentages of mice that experienced SRS, and open white bars represent percentages of mice that were SRS-free, during each treatment period. Differences between vehicle and PHT groups were not significant (ns, Fisher’s Exact Test). Likewise, differences between vehicle and CBZ groups were not significant (ns, Fisher’s Exact Test).

#### 3.3.2. Carbamazepine

Figure 6 (panels E-H) illustrates the effect of another sodium channel inhibitor, CBZ, on SRSs under our experimental conditions. All analyses were performed as described above for PHT. The average number of daily SRSs per mouse over the baseline period ranged from 0.7±0.2 to 1.3±0.4 SRSs/day (Figure 6, panels E-F); the weekly average was 1.0±0.2 SRSs/day (Figure 6G, baseline). There was no significant difference between the VEH (0.6±0.1 SRSs/day) and CBZ (0.4±0.1 SRSs/day) 5-day treatment averages (P = 0.1466, Wilcoxon Test, Figure 6G). Finally, as illustrated in Figure 6H, 7 of 22 mice were seizure-free during VEH treatment and 8 of 21 during CBZ treatment; this difference was not significant (P > 0.9999, Fisher’s Exact test).

#### 3.3.3. Valproic Acid

Figure 7 (panels A-D) illustrates the effect of VPA, a mixed-mechanism ASD, on SRSs under our experimental conditions. The average number of daily SRSs per mouse over the baseline period ranged from 1.1±0.6 to 2.6±1.3 SRSs/day (Figure 7, panels A-B); the weekly average was 2.0±0.8 SRSs/day (Figure 7C, baseline). The difference between the VEH (1.2±0.4 SRSs/day) and VPA (0.4±0.3 SRSs/day) 5-day treatment averages was significant (P = 0.0095, Wilcoxon Test, Figure 7C). Finally, as illustrated in Figure 7D, 1 of 13 mice was seizure-free during VEH treatment and 6 of 13 during VPA treatment; this difference was not significant (P = 0.0730, Fisher’s Exact test).

**Figure 7:**
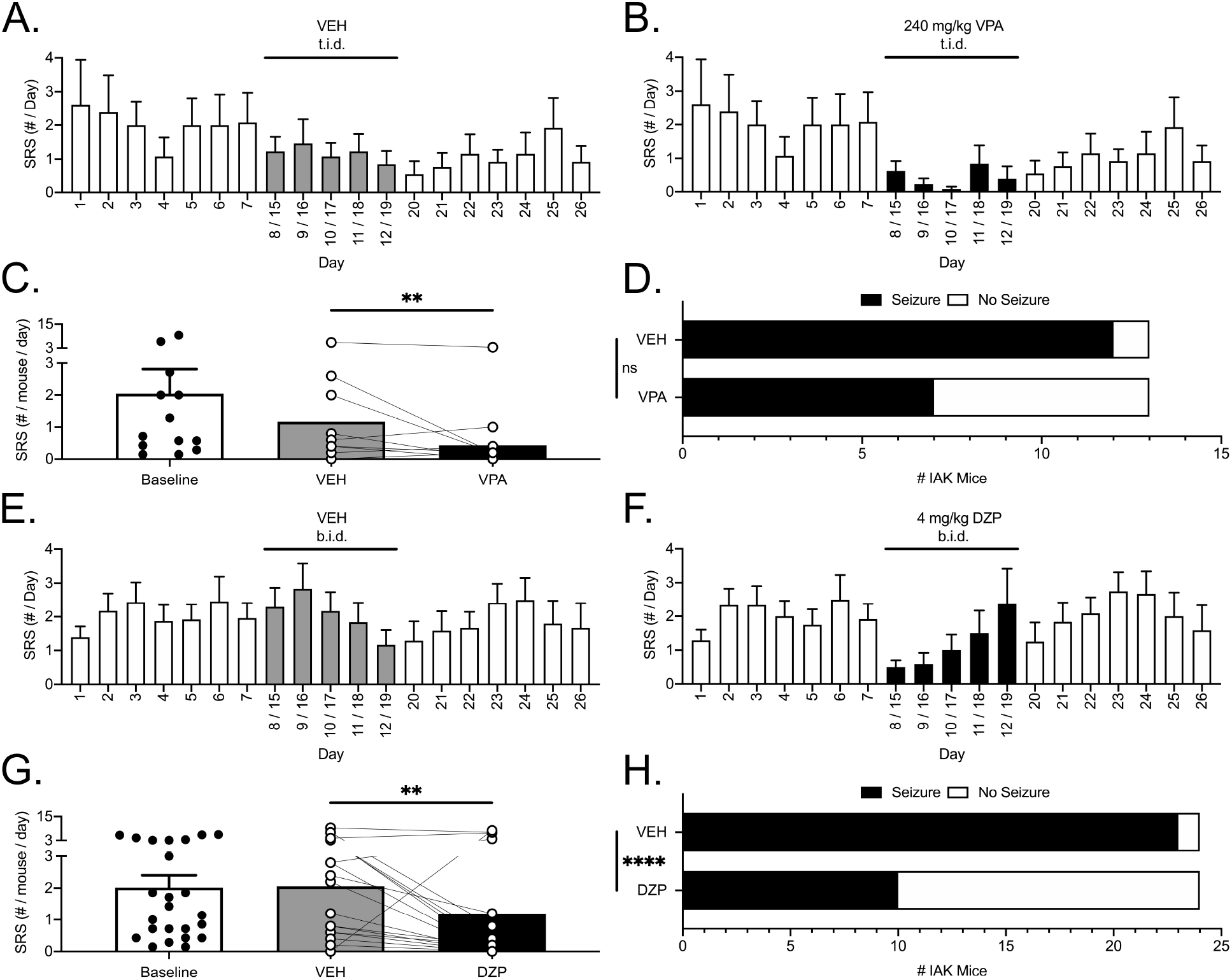
SRSs in IAK mice are sensitive to valproic acid (VPA) (720 mg/kg/day) and diazepam (DZP) (8 mg/kg/day). A-B and E-F) Daily average SRS frequencies over 26-day evaluation periods for vehicle (A: saline and E: saline) and (B) VPA or (F) DZP treatments. In all panels, open bars represent baseline and washout periods, while gray bars represent vehicle treatment averages (VEH) and solid black bars represent VPA or DZP treatment averages for both halves of the crossover. C and G) SRS frequency daily averages, and individual mouse data points, for baseline (open bar), vehicle (gray bar), and (C) VPA or (G) DZP (black bar) treatments. Vehicle and VPA data were significantly different (**, Wilcoxon Test). Likewise, Vehicle and DZP data were significantly different (**, Wilcoxon Test). D and H) Comparisons of the percentages of SRS-free mice during vehicle and (D) VPA or (H) DZP treatments. Solid black bars represent percentages of mice that experienced SRS, and open white bars represent percentages of mice that were SRS-free, during each treatment period. Differences between vehicle and VPA groups were not significant (ns, Fisher’s Exact Test). However, differences between vehicle and DZP groups were significant (****, Fisher’s Exact Test).

#### 3.3.4. Diazepam

Figure 7 (panels E-H) illustrates the effect of DZP, a GABA-A receptor modulating ASD, on SRSs under our experimental conditions. The average number of daily SRSs per mouse over the baseline period ranged from 1.3±0.3 to 2.5±0.7 SRSs/day (Figure 7, panels E-F); the weekly average was 2.0±0.4 SRSs/day (Figure 7G, baseline). The difference between the VEH (2.1±0.5 SRSs/day) and DZP (1.2±0.5 SRSs/day) 5-day treatment averages was significant (P = 0.0051, Wilcoxon Test, Figure 7G). Finally, as illustrated in Figure 7H, 1 of 24 mice was seizure-free during VEH treatment and 14 of 24 during DZP treatment; this difference in seizure freedom was significant (P < 0.0001, Fisher’s Exact test).

#### 3.3.5. Phenobarbital

Two doses of phenobarbital (PB) were tested. Figure 8 (panels A-D) illustrates the effect of 25 mg/kg b.i.d. PB, a barbiturate class GABA-A receptor modulating ASD, on SRSs under our experimental conditions. The average number of daily SRSs per mouse over the baseline period ranged from 1.1±0.4 to 2.0±1.1 SRSs/day (Figure 8, panels A-B); the weekly average was 1.5±0.4 SRSs/day (Figure 8C, baseline). The difference between the VEH (0.9±0.3 SRSs/day) and 25 mg/kg PB (0.4±0.2 SRSs/day) 5-day treatment averages was significant (P = 0.0134, Wilcoxon Test, Figure 8C). As illustrated in Figure 8D, 10 of 29 mice were seizure-free during VEH treatment and 15 of 29 during 25 mg/kg PB treatment; this difference was not significant (P = 0.2289, Fisher’s Exact test). Figure 8 (panels E-H) also illustrates the effect of 50 mg/kg b.i.d PB. The average number of daily SRSs per mouse over the baseline period ranged from 1.6±0.6 to 2.8±1.1 SRSs/day (Figure 8, panels E-F); the weekly average was 2.0±0.7 SRSs/day (Figure 8G, baseline). The difference between the VEH (1.9±0.7 SRSs/day) and 50 mg/kg PB (1.0±0.4 SRSs/day) 5-day treatment averages was significant (P = 0.0405, Wilcoxon Test, Figure 8G). Finally, as illustrated in Figure 8H, 4 of 18 mice were seizure-free during VEH treatment and 8 of 18 during PB treatment; this difference in seizure freedom was not significant (P = 0.2890, Fisher’s Exact test).

**Figure 8:**
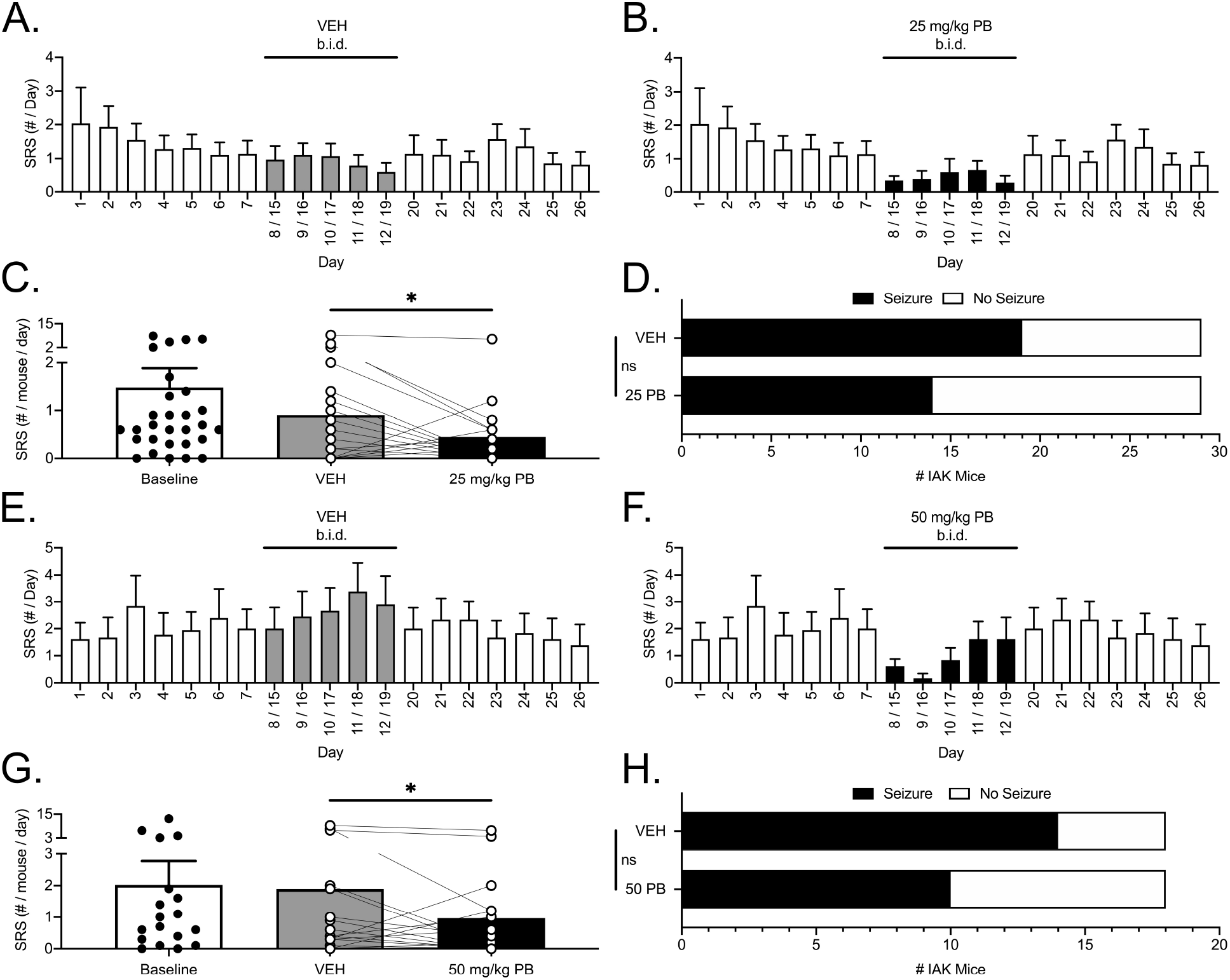
SRSs in IAK mice are sensitive to phenobarbital (PB) (50 mg/kg/day and 100 mg/kg/day). A-B and E-F) Daily average SRS frequencies over 26-day evaluation periods for vehicle (A and E: saline) and (B) 25 mg/kg b.i.d. or (F) 50 mg/kg b.i.d. PB treatments. In all panels, open bars represent baseline and washout periods, while gray bars represent vehicle treatment averages (VEH), and solid black bars represent 25 or 50 mg/kg b.i.d. PB treatment averages for both halves of the crossover. C and G) SRS frequency daily averages, and individual mouse data points, for baseline (open bar), vehicle (gray bar), and (C) 25 mg/kg b.i.d. or (G) 50 mg/kg b.i.d. PB (black bar) treatments. For both doses, vehicle and PB data were significantly different (*, Wilcoxon Test). D and H) Comparisons of the percentages of SRS-free mice during vehicle and (D) 25 mg/kg b.i.d. or (H) 50 mg/kg b.i.d. PB treatments. Solid black bars represent percentages of mice that experienced SRS, and open white bars represent percentages of mice that were SRS-free, during each treatment period. Differences between vehicle and PB groups, for either dose, were not significant (ns, Fisher’s Exact Test).

## 4. Discussion

The objective of the studies presented was to evaluate a mouse model of post-SE acquired MTLE based on microinjection of KA into the BLA. Although the most essential element of this model, microinjection of KA into the BLA, has been used by a number of laboratories for a variety of purposes, several methodological variations have been reported. In evaluating the effects of some of these variables, our purpose was not to resolve any disparity between our data and the data reported by others; instead, our goal was to develop a version of this model that maintained most of the essential experimental elements of these models described elsewhere, while optimizing quantitative endpoints suitable for future studies on the pharmacology of refractory seizures.

Beginning with the early rat studies of this model (Ben-Ari et al., 1980, 1979), use of a benzodiazepine to attenuate SE and improve survival is commonly reported. However, mortality data supporting this assertion in variants of the IAK mouse model are, to our knowledge, limited. Early model establishment work performed in the Henshall laboratory reported <10% mortality (presumably in the first 24 hours after SE) when SE is left uninterrupted; conversely, all mice that were given diazepam 40 min after initiation of SE survived (Araki et al., 2002). Similar implementations of this model that administered a single benzodiazepine, or intermittent benzodiazepine injections over a longer time period, have been shown to attenuate SE and result in low mortality over the first 24 hours (<5%) or over the first 3 days (7.5%) (Iori et al., 2016; Liu et al., 2013). In contrast, another reported model variant that did not administer a benzodiazepine reported 44% mortality (Welzel et al., 2020). Data for mortality observed over longer time periods after SE is similarly sparce. One report indicated 37.5% perish over the first 2 weeks post-SE (Kondratiuk et al., 2015), while another report indicated 8% of their mice died over a 3-day observation period that occurred 4 weeks post-SE (Engel et al., 2013). To our knowledge, the data described in this report is the first to directly compare IAK mice administered a benzodiazepine (midazolam) to a separate cohort that were allowed to experience uninterrupted SE. Furthermore, this study is also the first to continuously monitor IAK mice over a 90-day period. Surprisingly, we did not observe a statistically different mortality percentage between these experimental groups. When data from several cohorts of IAK mice that experienced uninterrupted SE were combined, mortality over the first 4 weeks and over the entire 90-day observational period was 20% and an additional 15% respectively. Although this additional 15% mortality over approximately 2 months was manageable for our subsequent pharmacology studies, methods that lower this rate and improve overall health of IAK mice in these experiments are being explored (e.g. use of wireless EEG telemetry implants versus traditional tethered methods that limit mouse mobility and self-care).

Another potential consequence of administering a benzodiazepine to attenuate SE is to limit the degree of resulting cellular damage. Previous work demonstrated that administering the benzodiazepine diazepam time-dependently attenuated contralateral hippocampal damage while preserving the relatively selective ipsilateral damage to the CA3 and CA1 subfields (Araki et al., 2002). However, it is noteworthy that, in this same report, hippocampal sub-regional damage was also examined when no benzodiazepine was administered; when SE induced with 3 µg of KA is uninterrupted by diazepam, the majority of hippocampal damage observed is still primarily observed in the ipsilateral CA3, followed to lesser degrees by ipsilateral CA1, contralateral CA3 and CA1, respectively. The data presented in the present study shows a similar pattern of damage regardless of having SE attenuated with the benzodiazepine midazolam. Considering that midazolam similarly did not affect mortality, the percentage of mice that developed SRSs, or the average daily frequencies of SRSs over a 90-day observation period, we elected to adopt a version of this model that doesn’t administer a benzodiazepine during SE.

Epilepsy, as defined as the occurrence of SRSs, was initially reported to develop with a latency of 2-5 days after KA-induced SE in IAK mice (Mouri et al., 2008). Subsequent experiments from a number of laboratories using subtle variants of this model have largely confirmed this latent period (Iori et al., 2016; Kondratiuk et al., 2015; Liu et al., 2013; Welzel et al., 2020). In the experiments reported here, it was impossible to definitively measure the latent period because video EEG was not recorded for the first 3 days after SE; early pilot experiments suggested that recovery from SE was improved when mice were allowed to freely move about their cages and care for themselves without the hindrances caused by an EEG tether. Since onset of epilepsy has been defined by others to be the occurrence of SRSs at least 48 hours from the end of SE (Iori et al., 2016), any SRSs that may have occurred over the first 48 hours should not influence the latent period by this definition. With consideration of this caveat, we did estimate the median latent period in our mice to be approximately 11 days (mean = 20 days). One possible reason for this discrepancy is that previous reports indicate all mice having SRSs within the relatively brief observation periods used in those studies (typically less than 2 weeks post SE). In our study, 30 of the 55 mice that developed SRSs did so with a latent period <2 week (mean 7.7±0.4 days). However, the remaining 25 mice exhibited their first SRSs between 15-74 days post-SE and had a correlated overall lower seizure frequency. While we anticipate that larger doses of KA may reduce the latency to first SRS and increase overall seizure frequency, we similarly expect an increased mortality under these experimental conditions. Accordingly, adjusting the dose of KA to optimize quantitative outcomes critical for specific experiments will continue to be an important consideration.

For the purposes of using the IAK mouse as a platform for testing the efficacy of investigational ASDs against pharmacoresistant seizures, two parameters expected to affect the suitability of the model are the relative stability of SRSs and their resistance to existing ASDs. Regarding the former, previous reports are laudatory of the model for its relatively stable electroclinical SRS frequencies in the range of approximately 0.5 – 5 SRSs per day, depending on the report (Frigerio et al., 2018; Iori et al., 2016; Jimenez-Pacheco et al., 2016; Kondratiuk et al., 2015; Liu et al., 2013; Mouri et al., 2008; Silva et al., 2016; Welzel et al., 2020). However, most of these reports evaluated SRS frequency for brief EEG recording periods (typically lasting 1-2 weeks) immediately after SE and/or approximately 2 months after SE. To our knowledge, the data presented here is the first to monitor SRSs continuously for 90 days in IAK mice. Several notable observations can be made regarding these data. First, these data show an apparent association between latent period and seizure frequency; specifically, earlier latency to the first SRS is associated with a higher average SRS frequency. A practical consequence of this association is that IAK mice diagnosed with epilepsy in the first 2 week after SE are subsequently enrolled in pharmacology studies and are simultaneously those with the highest SRS frequencies. Additionally, these same IAK mice with latent periods less than 14 days experienced progressively more frequent SRSs over the 90-day observation period. Similar progressive increases in SRS frequency were also observed in previous studies from different laboratories (Iori et al., 2016; Liu et al., 2013). Specifically, Iori et al. measured an average 3-fold progressive increase in SRS frequency over 2.5 months from the onset of epilepsy; the data presented in this report similarly shows a 3.75-fold increase in SRS frequency between week 1 and 9 after SE. Finally, and in contrast with these reported increases in SRS frequency, a different study indicated that SRS frequency did not appear to progress over 2 months; instead, SRS frequency trended to decline by approximately 3-fold (Welzel et al., 2020). While not significant, there was a trend in the data presented here for decreasing SRS frequencies between weeks 9 and 13.

Taken together, these progressive increases (and possibly decreases) in average SRS frequency appear to occur over a slow enough time period to permit experimental pharmacology studies that require 3-5 weeks to complete. However, another variable that could potentially confound pharmacology experiments is seizure clustering (Haut, 2006). While there is no universally accepted definition of seizure clusters, generally speaking, they can be defined as a closely grouped series of SRSs that are separated by a finite SRS-free period of time. As for specifics, numerous definitions of seizure clusters have been proposed and used (Bortel et al., 2010; Goffin et al., 2007; Hosford et al., 2016; Lim et al., 2018; Williams et al., 2009). The data presented here were analyzed for seizure clustering according to the definition proposed by Lim et al. (2018): the occurrence of one or more seizures per day for at least three consecutive days and at least five seizures within the cluster period. By this definition, SRS clusters in this data set lasted an average of 7 days and contained an average of 24 SRSs per cluster. However, as it pertains to determining the effects of investigational ASDs on SRSs, the duration of SRS-free periods is expected to have a greater impact on study design since the coincidence of SRS-free periods with drug exposure could result in a false-positive drug effect. The median SRS-free duration measured in the present data was 4 days. Since the pharmacology experiments described here utilized a 5-day crossover design, some of the IAK mice that were SRS-free during vehicle treatment (i.e. false positive effect) might be explained by this phenomenon.

As noted earlier, one of the primary objectives of this work was to evaluate this model of MTLE under the auspices of the ETSP and as part of its ongoing efforts to internally validate and adopt animal models of human epilepsy with pharmacoresistant SRSs. An important step towards this objective is to assess the sensitivities or resistances of SRSs in this model to a panel of approved and clinically used ASDs. This approach has been used for a number of animal models by the ETSP to characterize their pharmacoresistances to benchmark ASDs that can then serve as comparators for novel investigational compounds (Metcalf et al., 2017; Thomson et al., 2020; West et al., 2018, 2014). As an initial foray towards this objective, five ASDs from different mechanistic classes were tested for their effects on SRSs. With the exception of CBZ, this is the first study to report the effects of these ASDs on SRS frequency and seizure freedom in any version of the IAK model. The data presented demonstrates that SRSs in this IAK model of MTLE may be pharmacoresistant to two representative sodium channel-inhibiting ASDs (PHT and CBZ), but not to GABA receptor modulating ASDs (DZP and PB) or a mixed-mechanism ASD (VPA).

With regards to CBZ, only one other study by the Löscher laboratory has *directly* examined its effects on SRSs in a version of the IAK mouse model (Welzel et al., 2020). While oral administration of CBZ in IAK mice was used by Iori et al. to examine its potential disease modifying effects, its direct effects on SRSs were not analyzed (Iori et al., 2016). In the Löscher laboratory study, CBZ was administered intraperitoneally three times/day (for 5 days) with 30 mg/kg at 7:00 am, 30 mg/kg at 1:00 pm, and 45 mg/kg at 7:00 pm. This dosing regimen was able to significantly suppress SRSs, although an apparent tolerance to CBZ developed on day 5 possibly due to rapid elimination of CBZ in mice (Löscher and Schmidt, 2006). Similarly, the effects of CBZ presented in the present report show an apparent loss of effect after multiple administrations. Unlike the report from Welzel et al., and possibly due in part to the development of pharmacokinetic tolerance, the overall CBZ-mediated attenuation of SRS frequency was not statistically significant in our study. Other possible reasons for this difference may lie in dosing regimen differences (our study used 30 mg/kg t.i.d. while the Löscher study substitute a high 45 mg/kg dose at 7 pm) or interlaboratory methodological differences previously discussed.

Considering the mission of the ETSP, the ultimate objective of these studies was to determine if the IAK mouse represents a model of pharmacoresistant SRSs that can be used to evaluate novel investigational ASDs. With this objective in mind, it is of great importance to not only establish that SRSs in IAK mice are refractory to more than two clinically used ASDs, but also to assess how ASD activities compare between the IAK mouse and other rodent models of TLE. At the dose chosen in these experiments, SRSs in IAK mice were refractory to PHT (40 mg/kg/day). This dose was chosen in part based on pilot experiments where higher doses resulted in noted toxicity in IAK mice. Our laboratory has shown that SRSs in the i.p. KA spontaneously seizing rat model of MTLE are also refractory to PHT at similar doses (20 mg/kg/day) (Thomson et al., 2020). However, not all rodent models of TLE have SRSs that are resistant to PHT. For example, SRSs resulting from pilocarpine-induced SE were shown to be highly sensitive to PHT, although at a high dose of 100 mg/kg/day (Leite and Cavalheiro, 1995). While SRSs in the IAK mouse and i.p. KA spontaneously seizing rat exhibited a shared resistance to PHT and contrasted with the pilocarpine-induced SE rat where PHT was efficacious, the opposite correlation between models was observed regarding sensitivities to VPA. As shown here, VPA (720 mg/kg/day) significantly attenuated SRS frequency in IAK mice. Similarly, VPA (600 mg/kg/day) also significantly attenuates SRSs in the pilocarpine-induced SE rat (Leite and Cavalheiro, 1995). However, the same dose of VPA (600 mg/kg/day) was ineffective in the aforementioned rat i.p. KA SRS model (Thomson et al., 2020). Thus, these data suggest that IAK mice have a unique pharmacological sensitivity profile compared with the i.p. KA and pilocarpine rat models of TLE.

One of the ASDs representing GABA mimetics that was tested in this report was the benzodiazepine diazepam. At 8 mg/kg/day, diazepam significantly attenuated SRS frequency and increased the number of SRS free mice. Although not a model of SRSs, hippocampal paroxysmal discharges (HPDs) in a mouse intra-hippocampal KA model of TLE were shown to be significantly suppressed by diazepam (1 mg/kg) (Duveau et al., 2016). Furthermore, while diazepam was not tested in the i.p. KA spontaneously seizing rat model, other benzodiazepines (clobazam and clonazepam) were tested and shown to similarly attenuate SRS frequency and improve seizure freedom in that model (Thomson et al., 2020). Taken together, benzodiazepines appear to be efficacious in a number of models of TLE. However, adverse side effects, addiction liability, and tolerance tend to limit their clinical use in humans. With regards to the latter, tolerance to prolonged treatment with benzodiazepines has long been appreciated (Löscher and Schmidt, 2006). Despite achieving an overall statistically significant effect in our study, a tolerance-like daily decrease in diazepam’s attenuation of SRS frequency was observed.

The final ASD tested in this report was the barbiturate phenobarbital. Unlike the other ASDs tested herein, phenobarbital was tested at two doses (25 and 50 mg/kg b.i.d.). While both of these doses significantly attenuated SRS frequency, the magnitude of this attenuation was nearly identical for both doses (25 mg/kg b.i.d. produced a 50% attenuation, and 50 mg/kg b.i.d. produced a 51% attenuation of SRS frequency). These data suggest that SRSs in IAK mice are only partially sensitive to these doses of phenobarbital. In addition to the apparent tolerance observed with repeated 50 mg/kg dosing, the nature of this pharmacoresistance may be due, in part, to differential responsiveness between different IAK mice. Several IAK mice, notably those with high baseline SRS frequencies, showed little to no attenuation of SRS frequency in response to PB treatment at either dose. Similarly, and despite achieving an overall statistically significant decreased SRS frequency, 3 of 24 IAK mice were unaffected by diazepam (as indicated by their sustained average seizure frequencies between 3-9 SRS/day). These observations are reminiscent of the phenobarbital responder versus non-responder phenomenon known to occur in Sprague–Dawley rats where SRSs occur as a result of SE induced by prolonged electrical stimulation of the basolateral amygdala (Bethmann et al., 2007; Brandt et al., 2004).

## 5. Conclusions

In summary, the studies described here have characterized the methods, essential outcomes, and initial pharmacological profile of a mouse IAK induced SE model of MTLE. Unlike most methods reported and utilized by other laboratories, this version of the model does not arrest SE with a benzodiazepine. Since administration of midazolam did not result in significantly different latent periods, SRS frequencies over a 90-day observation period, or hippocampal damage 48 hours after SE, this step was determined to be unnecessary. Furthermore, reducing SE severity with a benzodiazepine may interfere with the efficacy of investigational treatments for epileptogenesis or disease modification (Welzel et al., 2020). Therefore, adoption of this methodological approach has the added benefit of developing these mice, and a foundational understanding of their disease progression characteristics, for future use in studies of investigational antiepileptogenic and/or disease modifying compounds. Finally, the sensitivities and resistances to ASDs reported herein provide an important first examination of the pharmacoresistance profile of this model. A more comprehensive examination of the effects of additional prototype ASDs, including additional dose response relationships combined with pharmacokinetic evaluations of plasma and brain concentrations, will be necessary in order to fully appraise the pharmacoresistance of SRSs in this model. However, the data presented here using five ASDs representing three mechanistic classes demonstrates that SRSs in these IAK mice are resistant to two prototype sodium channel inhibiting ASDs while being partially sensitive to GABA modulating ASDs and a mixed-mechanism ASD; these data alone make a compelling case for the designation of SRSs in these mice as pharmacoresistant. Accordingly, this model is being incorporated into the NINDS-funded ETSP testing platform for treatment resistant epilepsy.

## Acknowledgements

The authors would like to acknowledge and thank the laboratories of Dr. David C. Henshall and Dr. Annamaria Vezzani for their helpful technical guidance as we began to establish this model in the ETSP. Furthermore, we thank all the members of the ETSP contract site at the University of Utah (formerly known as the Anticonvulsant Drug Development, ADD, Program) for scientific, technical, and logistical support. In particular, we thank Kristina Johnson, Elisa Koehler, Gaëlle Batot, Dorina Diekjürgen, and Sharon Edwards. The authors would also like to thank Dr. Brian Klein and the ETSP at the National Institutes of Neurological Diseases and Stroke (NINDS) for their review and comments on this manuscript. Finally, we would like to acknowledge and thank the external consultation board members for the ETSP (Dr. Amy Brooks Kayal, Dr. Henrik Klitgaard, Dr. Wolfgang Löscher, Dr. James O. McNamara, and Dr. Jennifer A. Kearney) for their continuing helpful guidance.

## Abbreviations

ASD: antiseizure drugs
BLA: basolateral amygdala
CBZ: carbamazepine
DZP: diazepam
EEG: Electroencephalography
ETSP: Epilepsy Therapy Screening Program
FJB: FluoroJade B
GABA: gamma-Aminobutyric acid
IAK: Intra-Amygdala Kainate
ILEA: International League Against Epilepsy
KA: kainic acid
LTG-R: lamotrigine resistant
MTLE: mesial temporal lobe epilepsy
NINDS: National Institute of Neurological Disorders and Stroke
PB: phenobarbital
PHT: phenytoin
SE: status epilepticus
SRS: spontaneous recurrent seizure
TLE: temporal lobe epilepsy
VEH: vehicle
VPA: valproic acid / valproate

